# Fast and Simple Tool for the Quantification of Biofilm-Embedded Cells Sub-Populations from Fluorescent Microscopic Images

**DOI:** 10.1101/181495

**Authors:** Mikhail I Bogachev, Vladimir Yu Volkov, Oleg A Markelov, Elena Yu Trizna, Diana R Baydamshina, Vladislav Melnikov, Regina R Murtazina, Pavel V Zelenikhin, Irshad S Sharafutdinov, Airat R Kayumov

## Abstract

Fluorescent staining is a common tool for both quantitative and qualitative assessment of pro- and eukaryotic cells sub-population fractions by using microscopy and flow cytometry. However, direct cell counting by flow cytometry is often limited, for example when working with cells rigidly adhered either to each other or to external surfaces like bacterial biofilms or adherent cell lines and tissue samples. An alternative approach is provided by using fluorescent microscopy and confocal laser scanning microscopy (CLSM), which enables the evaluation of fractions of cells subpopulations in a given sample. For the quantitative assessment of cell fractions in microphotographs, we suggest a simple two-step algorithm that combines single cells selection and the statistical analysis. To facilitate the first step, we suggest a simple procedure that supports finding the balance between the detection threshold and the typical size of single cells based on objective cell size distribution analysis. Based on a series of experimental measurements performed on bacterial and eukaryotic cells under various conditions, we show explicitly that the suggested approach effectively accounts for the fractions of different cell sub-populations (like the live/dead staining in our samples) in all studied cases that are in good agreement with manual cell counting on microphotographs and flow cytometry data. This algorithm is implemented as a simple software tool that includes an intuitive and user-friendly graphical interface for the initial adjustment of algorithm parameters to the microphotographs analysis as well as for the sequential analysis of homogeneous series of similar microscopic images without further user intervention. The software tool entitled *BioFilmAnalyzer* is freely available online at https://bitbucket.org/rogex/biofilmanalyzer/downloads/.

## Introduction

One of the key issues in both pro- and eukaryotic cell studies is the quantitative characterization of cellular subpopulations like the estimation of the fractions of either live or dead cells in a given population, differentiation of bacterial species in mixed biofilms or eukaryotic cell types in culture. There are two common experimental approaches to these issues, namely the flow cytometry and the fluorescent microscopy. In both methods the cells are stained with fluorescent dyes which specifically differentiate the cells of interest. Thus, Syto9/PI, DioC6/PI, AO/PI, CFDA/PI, Calcein AM/PI, Hoechst/PI and many other combinations of dual staining are widely used to differentiate viable and non-viable cells [1–3]. Normally, the first dye is biochemically modified by viable cells followed by the production of the green-fluorescent product. The second dye like the propidium iodide or ethidium bromide penetrates through the damaged membrane of dead cells forming complexes with nucleic acids and providing red fluorescence. Despite of multiple reports that the estimation of viable cells fraction by using vital staining often exhibits significant differences in comparison with the values obtained by using classical microbiological methods [4], fluorescent staining remains a fast and easy approach to the quantification of (non-)viable cells.

While flow cytometry typically provides with a more accurate assessment of the cell subpopulation fractions [5, 6], it has several principal limitations that significantly narrow its application area [7]. In particular, cells being adhered to each other and to external surfaces should be suspended prior to their infusion into a cytometer that appears difficult when, for example, bacterial biofilms or strongly adherent cells are analyzed, or the original structure of the cell colonies, cell complex or tissue structure should be preserved. Moreover, flow cytometer is normally unable to detect particles <0.500 μm [8]. Finally, currently available flow cytometry systems require considerable amount of maintenance and highly skilled operators.

Fluorescent microscopy is largely free of above limitations and provides a reasonable alternative to the cytometric measurements. However, in the presence of adherent and/or spore-like cells they largely overlap leading to the limitations of direct cell selection and counting algorithms in the microscopic images. The situation gets even more complicated when the cells are not equidistantly stained, image quality and color balance varies in different fields of view. Despite of the above limitations, manual counting is usually still possible, while it requires significant efforts from experts increasing the lab personnel workload drastically. Thus, automatic or semi-automatic analysis of cells seems to be a fast and easy approach to the microscopic data quantification. In the last two decades, a number of methods and computer-assisted algorithms have been developed to resolve the cell counting issue implemented in a number of both commercial and free software tools [9–13]. Existing software solutions include cell counting and classification algorithms [14], estimation of their parameters from microscopic imaging [15], 3d reconstructions from confocal microscopy data [16] and several other more specific applications. However, automatic microscopic image analysis remains challenging in the presence of adherent and/or spore-like cells that are common conditions in biofilm studies. Automatic counting methods are usually based either (i) on detection, selection and counting of discrete objects, or (ii) on the statistical analysis of the image properties that avoid direct counting approach and estimate some effective characteristics from the statistical properties of the entire image [17]. While the detection methods fail under cell overlapping conditions, the statistical assessment methods are unable to differentiate between various types of cells. Among few exceptions that largely overcome the above limitations, a very recently designed software tool for the quantification of live/dead cells in a biofilm based on a series of image transformation could be mentioned [18].

Another common disadvantage of many automatic image analysis tools in practical settings is often, though may sound surprising, their excessive use of automation. Quite often image analysis algorithms use complex transformations and filtering procedures with parameters that can be hardly controlled by the user who is often not an expert in digital image processing. As a result, there is little or no feedback between the algorithm and its end user. Thus, the user deals with a kind of black box design, where an image is inserted and a value comes out, without being able to cross-check the performance of the algorithm at some intermediate steps. Despite the increasing complexity and performance of image processing tools, the variety of cell structures and microscopic imaging conditions to our opinion is still too broad to fully rely upon automation in all cases.

Here we suggest an algorithm based on a simple combination of the object counting and statistical approaches implemented in an easy-to-use cell-counting software tool, *BioFilmAnalyzer*, freely available at https://bitbucket.org/rogex/biofilmanalyzer/downloads/. Following preliminary threshold-based filtering and segmentation of the image, an effective number of cells is calculated under partially-manual control by the investigator. To support the investigator with finding the right balance between the parameters that should be adjusted at the initial segmentation step, namely the selection threshold and the typical size range of single cells, we suggest a simple procedure that includes estimation of the histograms of the sizes of isolated objects after threshold-based filtering and an objective criterion based on the analysis of these histograms. Based on a series of experimental measurements performed in bacterial cells of *S. aureus* and *B. subtilis* exhibiting different shapes as well as eukaryotic cells, we show explicitly that the suggested approach effectively accounts for the fractions of live/dead cells in all studied cases. The validity of the *BioFilmAnalyzer* based cells live/dead fractions quantification was assessed by comparison with the results of manual counting performed by several experts in visual microscopic image analysis and cytometric measurements.

## Materials and methods

### Bacterial strains, cell lines and fluorescent microscopy

The fluorescent microscopic images of bacterial cells obtained in previous works were used [19–23]. Briefly, *Staphylococcus aureus* (ATCC^®^ 29213™) and *Bacillus subtilis 168* grown in 35-mm TC-treated polystyrol plates (Eppendorf) for 48 hours under static conditions at 37°C to obtain rigid biofilm structures were further subjected to differential live/dead fluorescent staining.

The human colon adenocarcinoma Caco-2 cells (RCCC) were cultured in DMEM supplemented with 10% FBS, 2 mM L-glutamine, 100 μg ml^-1^ penicillin and 100 μg ml^-1^ streptomycin. The cells were seeded in 24-well plates at the density of 30000 cells per well and allowed to attach overnight. The cells were cultured at 37 °C and 5% CO_2_ until 70% confluence and camptothecin (Sigma-Aldrich) was added in final concentration of 6 μM. After 24 h of exposition the cells were subjected to fluorescent staining and analyzed with flow cytometry and fluorescent microscopy.

The viability of biofilm-embedded cells was evaluated by staining for 5 min with the Acridine orange (Sigma) at final concentration of 0.12 μg/ml (green fluorescence) or 3,3’-Dihexyloxacarbocyanine iodide (DioC6) (Sigma) at final concentration of 0.02 μg/ml (green fluorescence) and propidium iodide (Sigma) at final concentration of 3 μg/ml (red fluorescence) to differentiate between bacteria with intact and damaged cell membranes (live and dead cells). The eukaryotic cells were stained with DioC6 (0.02 μg/ml) and propidium iodide (3 μg/ml). The microscopic imaging was performed using either Carl Zeiss Observer 1.0 microscope or Carl Zeiss CLSM780 with 40-100× magnification.

### Flow cytometry

Cytometric analysis was performed using BD FACSCanto II flow cytometer. The Caco-2 cells were stained as described above. Following data analysis was carried out using FACSDiva software.

### Image analysis methodology

Image analysis starts with the preliminary detection of pixels with given color channel intensity (red, green, blue, or overall) exceeding a given threshold. Fig 1A shows a sample of original microscopic image, while Fig 1B shows the result of threshold-based selection of the red channel according to the differential rule *I_R_* – *I_G_* > *T*, where *I_R_* and *I_G_* are the intensities in the red and in the green channels, respectively, *T* = 30 is the threshold setting. In the next step, the preprocessed image is segmented into separate non-overlapping objects by adjusting horizontally, vertically or diagonally neighboring pixels above the threshold to the same object. Additionally, objects with a given set of properties that determine the cell subpopulation are selected. Fig 1C exemplifies the results of selection by size, with objects only in a given size range between *s*_min_ = 500 and *s*_max_ = 1000 pixels being highlighted, where the color encodes each object.

**Fig 1.**
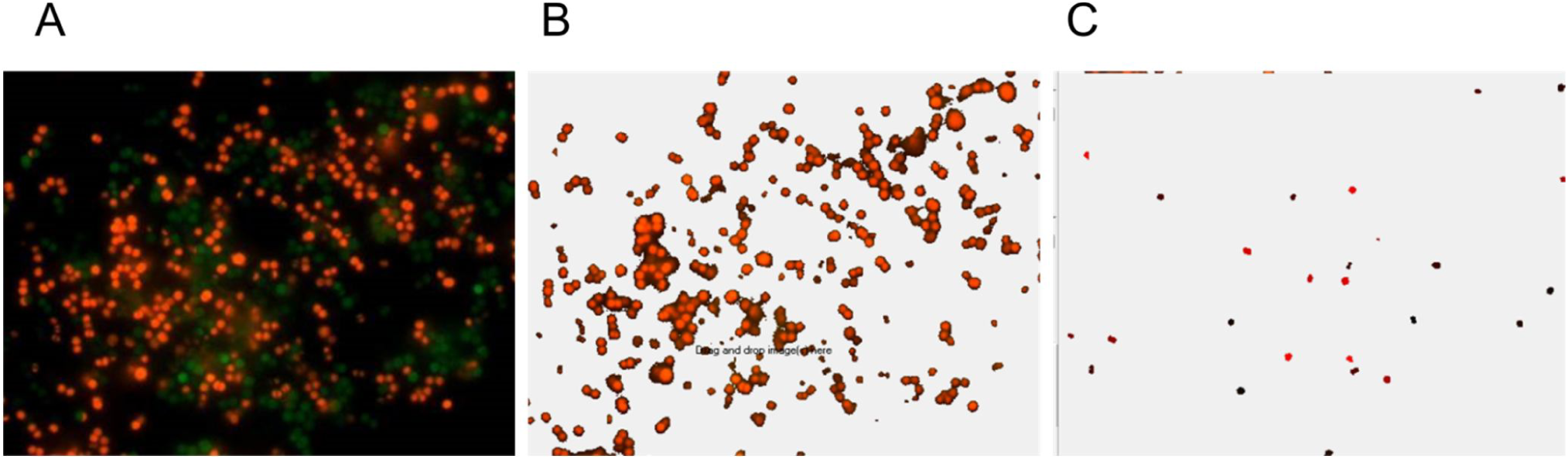
Consecutive steps in cell quantification by *BioFilmAnalyzer*. (A) original image containing overlapped red and green channels, (B) selected red channel data after threshold-based filtering, (C) selected cells of size between *s*_min_ and *s_max_* that are used to determine the effective single cell size with each separate cell shown by another color as determined by the automatic segmentation algorithm.

In the following, we count the selected objects, calculate their total area, and obtain the average size of the typical cell from the studied subpopulation. Next we determine the effective number of cells by dividing the total area above the threshold (Fig 1B) by the average size of the selected cells (Fig 1C) according to

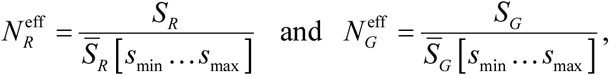

where 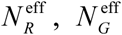 are the effective numbers of cells in the red and the green channels, respectively; *S_R_, S_G_* are the total area of selected cells belonging to the red and the green channels, respectively; 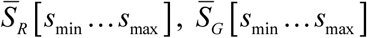 are the average area of cells with a size range between *s*_min_ and *s*_max_.

While *T, s*_min_ and *s*_max_ have to be appropriately chosen, their choice has to be performed once in a series of similar experiments considering similar cell types as well as similar staining and imaging conditions. Accordingly, so far their manual choice by expert seems to be the easiest solution option, since the feedback from the samples tested first allows for a more specific adjustment of these parameters, to avoid potentially bizarre results that may arise in the case of blind choice of algorithm parameters with no feedback. In the following, for the entire image series the parameters are fixed and no further manual adjustment is required. Thus a series of images representing different fields of view under identical conditions in simply passed through an algorithm. When the differences in the conditions are minor and do not change significantly the microscopic images, but influence only some of their parameters like the live/dead cell fractions, such as testing antimicrobials with gradually changing concentrations, several series of images can be submitted without further adjustment of the algorithm parameters. The implemented software solution organizes the results of calculations in a table that could be exported and represents them graphically.

### Statistical assessment

Here we used simple linear regression without the intercept term, i.e. *y* = *kx*. In the suggested regression model the ideal case corresponds to *k* = 1 or simply *y* = *x* that would mean perfect agreement between automatic and manual expert counting. The quality of the results is characterized by two independent coefficients. The first one is the standard coefficient of determination 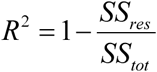, where *SS_tot_* is the total sum of squares 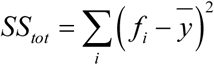 and *SS_res_* is the residual sum of squares 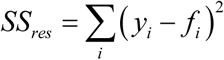, here 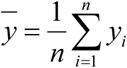 is the mean value of all analyzed data points and *f_i_* is calculated regression point. The *R*^2^ coefficient indicates how well the analyzed data set is replicated by the regression model.

In our study it was also important how close is the regression model to the ideal case *y* = *x*. Therefore we introduced a similarly designed metric of how well the model *y* = *kx* is close to the ideal counting line *y* = *x* which we denote here by *L*^2^. Its definition is similar to *R*^2^ besides the calculation is performed for the regression points *f_i_. SS_res_* for *L*^2^ is calculated as 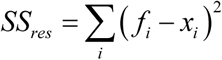, where *x_i_* is the corresponding abscissa value for the current *f_i_*, and *SS_tot_* is the same as in *R^2^*. In ideal case when observational regression line follows 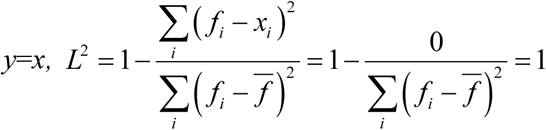. Thus both *R*^2^ and *L*^2^ coefficients range from 0 to 1.

## Results and Discussion

### Software description

*BioFilmAnalyzer* software (S1 and S2 Files) can be used to count any bacterial or eukaryotic fluorescent stained cells from two-dimensional microphotographs provided either as single images or as series of images obtained under similar conditions such as staining, magnification and color balance (for example, several views of the same sample taken with fixed camera settings). This in-house algorithm was implemented in C++ programming language and is compatible with Windows (2000/XP versions and higher). The software tool has a simple user-friendly environment for the image analysis with the two-step algorithm allowing manual parameter adjustments by end user at the initial step and automatic image series analysis at the second step. The logic of the image analysis is based on the preliminary adjustment of the algorithm parameters by using one or several images that the investigator finds more or less representative for the studied cohort in terms of imaging conditions. In the first step, simple drag & drop of a single image into the program window initiates its instant processing including threshold based detection of cells according to a specified rule based on the exceedance of a given threshold by either a certain color channel, difference between color channels or overall intensity. The threshold value *T* which is the only algorithm parameter in this first step can next be adjusted by the investigator by its increasing in cases of strongly autofluorescent background or by its decreasing in low contrast images until the background noise is eliminated. The view can be easily switched between the original and the processed images by a single click on the upper image panel.

For image series exhibiting strongly non-homogeneous color distributions in the studied color channels, there are two common solutions, including (I) adaptive adjustment of the threshold value *T* for each of the images individually, as well as (II) preliminary image color normalization that often appears a faster alternative. Since color normalization is a simple and standard image preparation procedure that is straightforward and thus can be applied consecutively to a series of images using many commercial or free image processing software tools (e.g., ImageMagick, Gimp etc.), we do not focus further on this issue. However, often significant image preprocessing leads to other problems, such as the increased level of the background noise that is enhanced together with the useful parts of the image when amplifying the intensity of the color channel with low intensity in the original image. For that reason, adaptive thresholding often appears a more efficient solution. As we show below, our algorithm is robust against moderate variations of the threshold *T* around its optimized value for a given color distribution, and thus the color normalization appears unnecessary for image series with moderate variations of color distributions. For those images exhibiting considerable variations of staining and imaging conditions, or pronounced color imbalance, the more preferable adaptive thresholding procedure is still possible.

In the second step, the effective cell size should be adjusted such that only or nearly only single cells appear in the lower image panel, which shows the processed image after a simple segmentation procedure. Increasing the lower limit of the cell size helps to eliminate some noise bursts, while decreasing the upper limit eliminates large patches of adherent cells from the effective cell size statistics. With the cell size window representing only a simple criteria that often appears insufficient, whenever necessary, further elimination of anomalous segments can be done manually by selecting and double-clicking over them in the lower image panel. Since the parameters including sizes of different sub-population of cells can differ from one another (e.g., non-viable eukaryotic cells are commonly smaller than viable cells), the effective cell size should be re-adjusted for each studied sub-population of cells. For this reason, the analysis of each color channel should be performed separately, and thus with the exception of the intensity based analysis rule, all other options analyze color channels individually. Thus they perform similarly for single or overlapped color channel images.

In general, the threshold *T* and single cell size range are interdependent. Since single cells are represented by inhomogeneous color intensity, with increasing the threshold *T* typical sizes of isolated objects representing single cells decrease, due to the removal of some pixels with intensity smaller than *T*. For that reason, it is important to keep the right balance between the chosen threshold *T* and the single cells size range between *s*_min_ and *s*_max_ In the following, we suggest an objective criterion for keeping such balance based on the analysis of the histograms of isolated objects sizes that is implemented in the software tool and validated by multi-threshold analysis.

Finally, the effective number of cells is determined as the total area of the image exceeding the threshold *T* shown in the upper panel divided by the effective cell size determined from the lower image panel. Once a reasonable set of algorithm parameters in found, a series of up to 100 images can be dropped onto the program window for fully automated analysis with the same set of parameters as determined from the first one or several representative images without further user intervention. Finally, the results can be exported to MS Excel for validation of the results and following statistical analysis. In comparison with some other freely available image processing and analysis software [18, 24, 25], our image analysis procedure is semi-automatic: the initial settings, while based on an objective statistical criterion, are modified by manual adjustment of the algorithm parameters and each step of the processing algorithm is fully visualized to enable expert controlled and assessment of the analysis quality. Once optimized for a representative sample image, the software tool next analyzes the entire image series automatically over coffee-break.

### Software validation by bacterial cells counting

We analyzed the efficiency of the proposed algorithm using the fluorescent images of coccal *(Staphylococcus aureus)* and rod *(Bacillus subtilis)* cell morphologies containing different fraction of viable (green-stained) cells. Since the cell suspension can be easily analyzed with flow cytometry, we focused on the analysis of adherent, biofilm-embedded cells. For that, we used previously obtained series of microphotographs of *S. aureus* in 48-72 h old biofilms treated with different antimicrobials [19, 21]. Cells were stained with DioC6 and propidium iodide to differentiate the viable and non-viable cells, and non-viable cells fraction was quantified with *BioFilmAnalyzer* software. As a representative example, twelve images with overlapped red and green channels containing different fraction of viable cells are shown in Fig 2. Since the image brightness, contrast and saturation vary from image to image depending on the staining quality, microscope settings and sample itself, for each microphotograph shown in Fig 2 the thresholds *T* (ranging from 25 to 55) and effective cell sizes were chosen individually. Next to validate the performance of the algorithm when the intensity threshold is chosen quite arbitrarily without careful manual adjustment to the imaging conditions, these microphotographs were analyzed consequently for different analysis thresholds (15, 20, 30, 45 and 60) and the results obtained by the automatic cell counting were plotted as a linear function *y* = *kx* of the manual cell counting (Figs 3A-3C). While the number of both red (non-viable, Fig 3A) and green (viable, Fig 3B) cells decreases at higher thresholds, their fractions (i.e., live/dead ratio) remained rather the same for each threshold value exhibiting no significant differences with the manual evaluation data (compare thick dashed line corresponding to the ideal fit of manual and automatic counting, Fig 3C).

**Fig 2.**
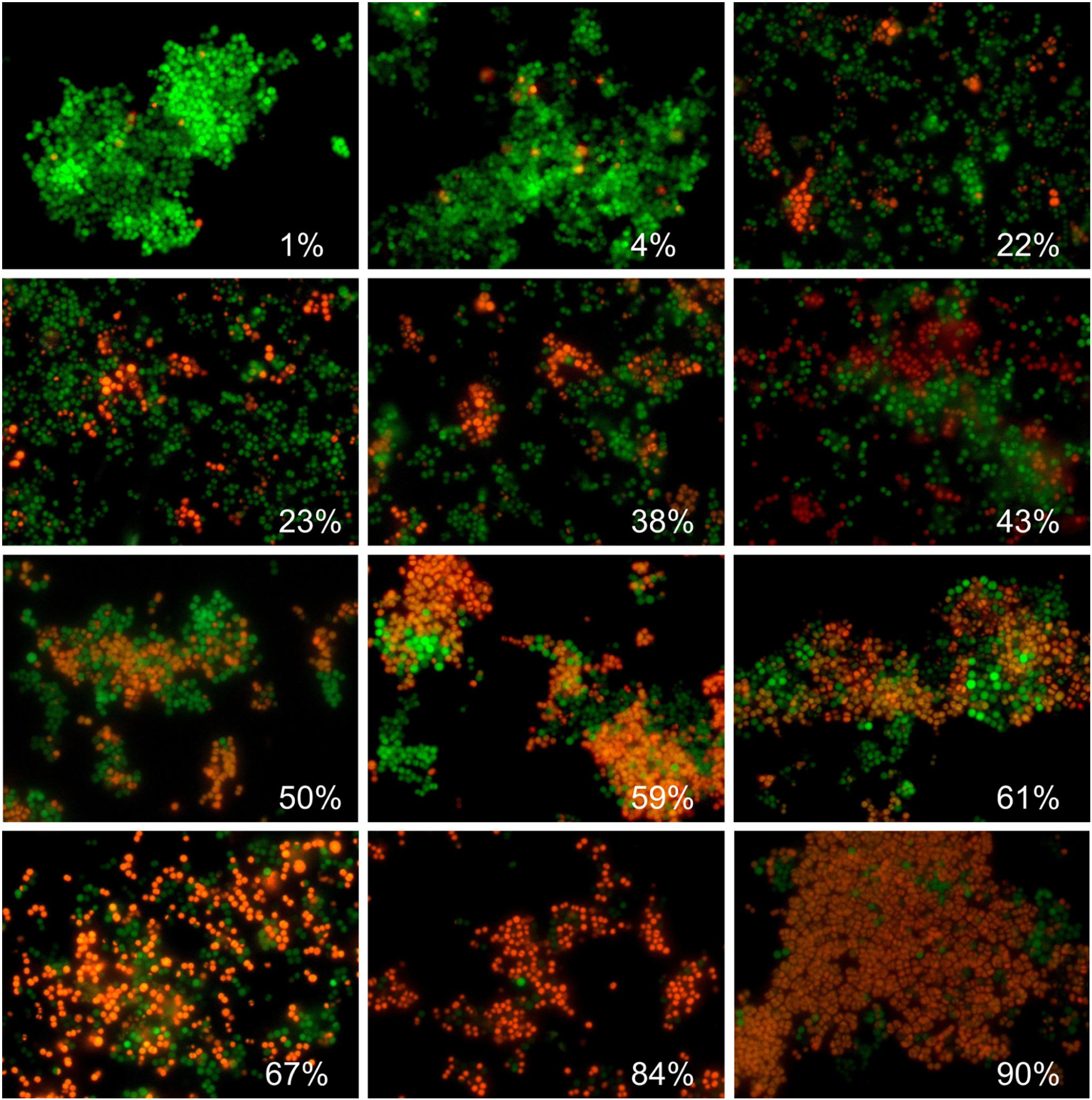
Evaluation of the *S. aureus* biofilm-embedded non-viable cells fraction by using *BioFilmAnalyzer* software exemplified for 12 microscopic images. The percentage of red-stained cells quantified by the *BioFilmAnalyzer* software is shown in each panel.

**Fig 3.**
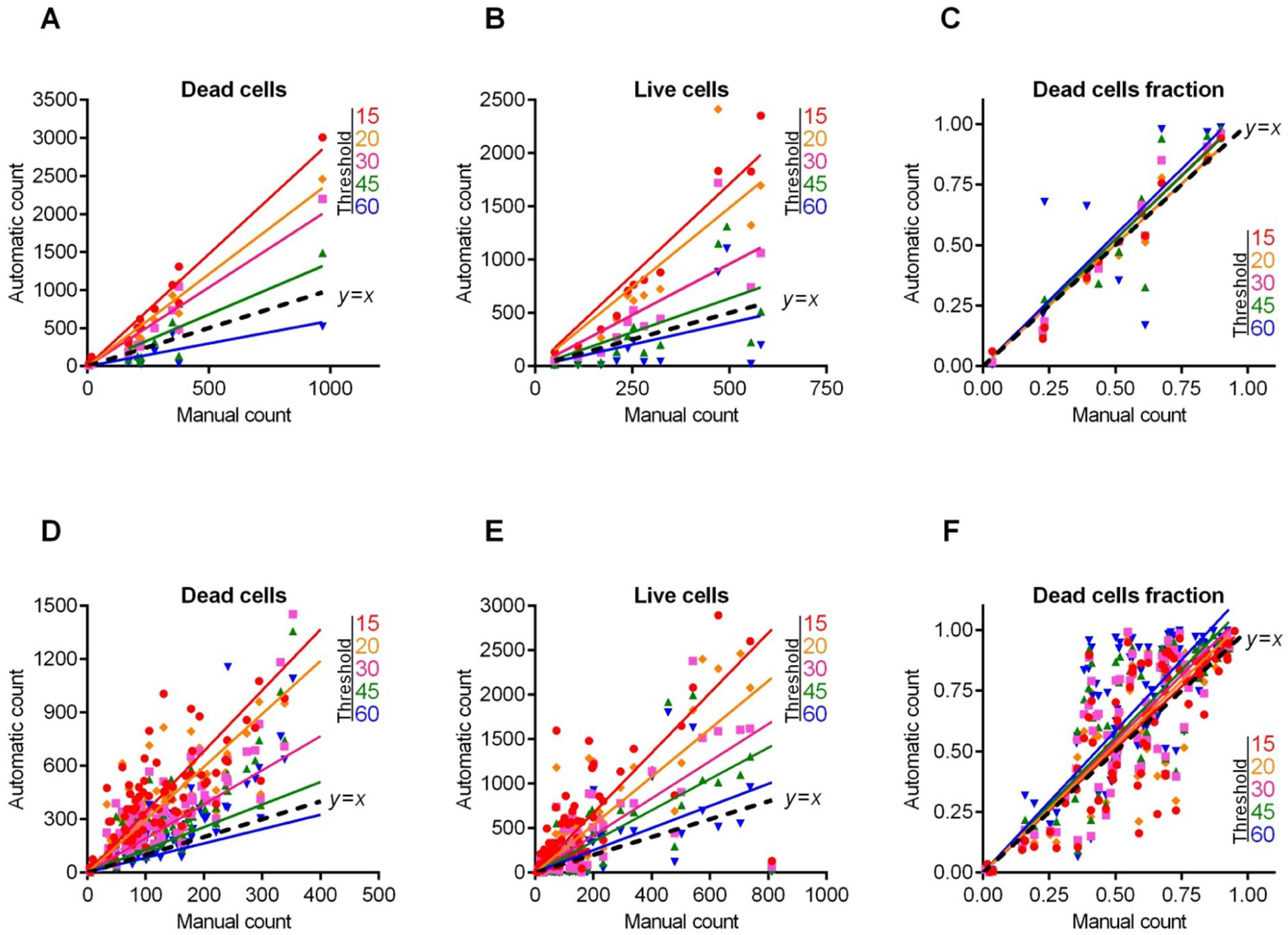
*S. aureus* cells count and live/dead ratio dependence on the image analysis threshold *T*. Full lines show the linear regression lines, while dashed line shows the ideal counting line as determined by the manual analysis performed by several experts in visual microscopic image analysis. Panels A-C show data for the 12 images presented in Fig 2. Panels D-F show data for the 115 microscopic images taken randomly from different experiments with various imaging conditions.

Table 1 shows the regression coefficients *k* for the linear regressions *y* = *kx* and their coefficients of determination R^2^. Since by definition there should be no systematic shift between the automatic and the manual count, i.e. in the absence of viable or non-viable cells the respective number of cells equals zero, we used the simplest linear regression model without intercept. Since the ideal counting corresponds to *k* = 1 or simply to the *y* = *x* line, we also calculated another coefficient denoted *L*^2^ that determines the deviation of the obtained regression line *y* = *kx* from the ideal counting line *y* = *x*. Of note, the automatic cell enumeration by *BioFilmAnalyzer* software exhibited the best fit with manual count at the analysis threshold of 45 (for images presented in Fig 2). For further details on the statistical analysis of our results, we refer to the Materials and Methods section.

**Table 1.**
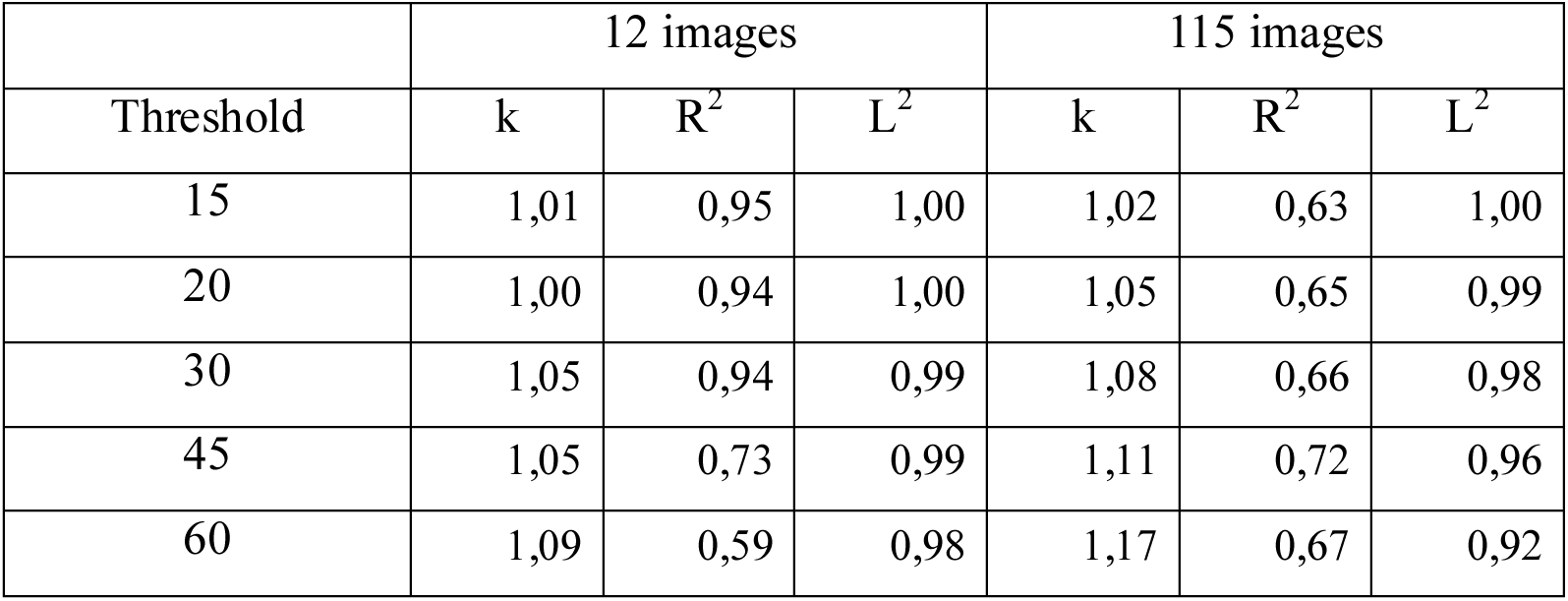
Regression coefficients *k*, the coefficients of determination *R*^2^ and the accuracy coefficient *L*^2^ indicating the correspondence between the automatic and the manual counting for the *S. aureus* live/dead ratios on fluorescent images.

|  | 12 images |  |  | 115 images |  |  |
| --- | --- | --- | --- | --- | --- | --- |
| Threshold | k | $R^2$ | $L^2$ | k | $R^2$ | $L^2$ |
| 15 | 1,01 | 0,95 | 1,00 | 1,02 | 0,63 | 1,00 |
| 20 | 1,00 | 0,94 | 1,00 | 1,05 | 0,65 | 0,99 |
| 30 | 1,05 | 0,94 | 0,99 | 1,08 | 0,66 | 0,98 |
| 45 | 1,05 | 0,73 | 0,99 | 1,11 | 0,72 | 0,96 |
| 60 | 1,09 | 0,59 | 0,98 | 1,17 | 0,67 | 0,92 |

Individual adjustment of the threshold and effective cell size is obviously possible only when a small number of images should be analyzed. For a more accurate quantification, a series of 10 or more images from the same sample should be analyzed. The *BioFilmAnalyzer* software allows analysis with constant settings of threshold and cell size of two or more images when they are dragged & dropped simultaneously onto the program window. To estimate the performance of the software when the images with different quality are analyzed without individual optimization of the intensity threshold, a series of 115 microphotographs randomly taken from different experiments were analyzed consequently for different thresholds (15, 20, 30, 45 and 60) with fixed cell size ranges used to determine the effective cell sizes. The results were compared against manual cell counting data (Figs 3D-3F). Similarly to previous results obtained for 12 images with accurate settings of both thresholds and cell sizes, the live/dead ratio remained almost the same for each threshold value and the obtained regression fit was in an excellent agreement with the manual cell counting data indicated by *L*^2^ being very close to 1.0 at thresholds in the range of 15-30 (Table 1).

Next, similar analysis was performed for the microscopic images of *B. subtilis* biofilm-embedded cells treated with different antimicrobials [20]. Like in the previous example, Fig 4 shows 12 representative images with different fraction of viable cells quantified with *BioFilmAnalyzer* software. Figs 5A–5C are designed similarly as Fig 3 and show the interdependence of the image analysis threshold and cell count calculated by *BioFilmAnalyzer*. Similar to *S. aureus* microphotographs, the cell count calculated automatically increased at high threshold with the best fit with manual count at threshold value of 45, while the live/dead ratio remained unchanged with *L*^2^ exceeding 0.96 at thresholds up to 45, suggesting that the performance of the algorithm does not depend on the fine tuning of the threshold and on the cell shape. Figs 5D-5F show the analysis of 50 randomly chosen images of live/dead stained *B. subtilis* cells. Similarly, statistical analysis did not reveal significant differences between automatic and manual live/dead ratio estimation (Table 2).

**Fig 4.**
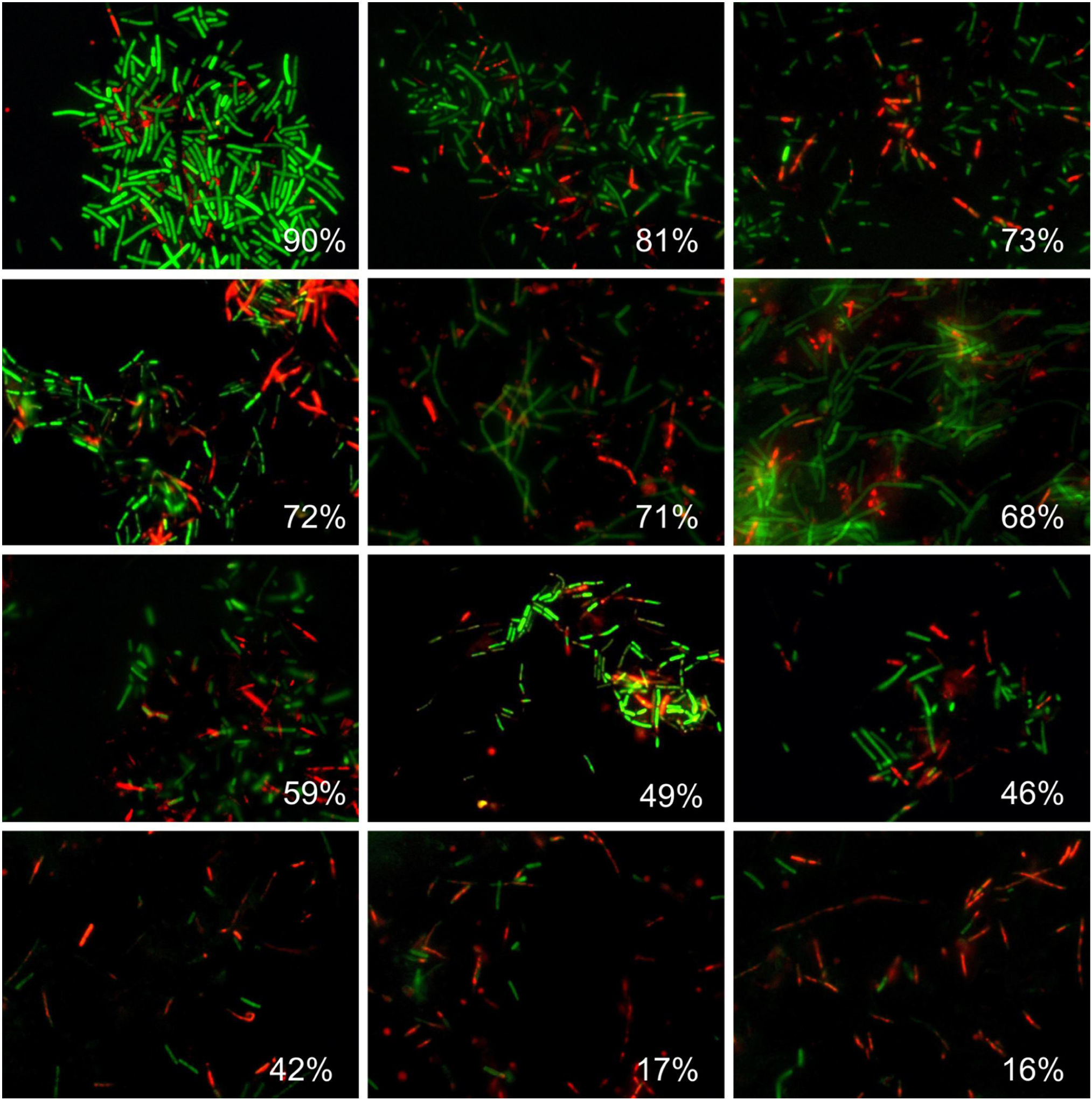
Evaluation of the *B. subtilis* biofilm-embedded non-viable cells fraction by using *BioFilmAnalyzer* software exemplified for 12 microscopic images. The percentage of red-stained cells quantified by the *BioFilmAnalyzer* software is shown in each panel.

**Fig 5.**
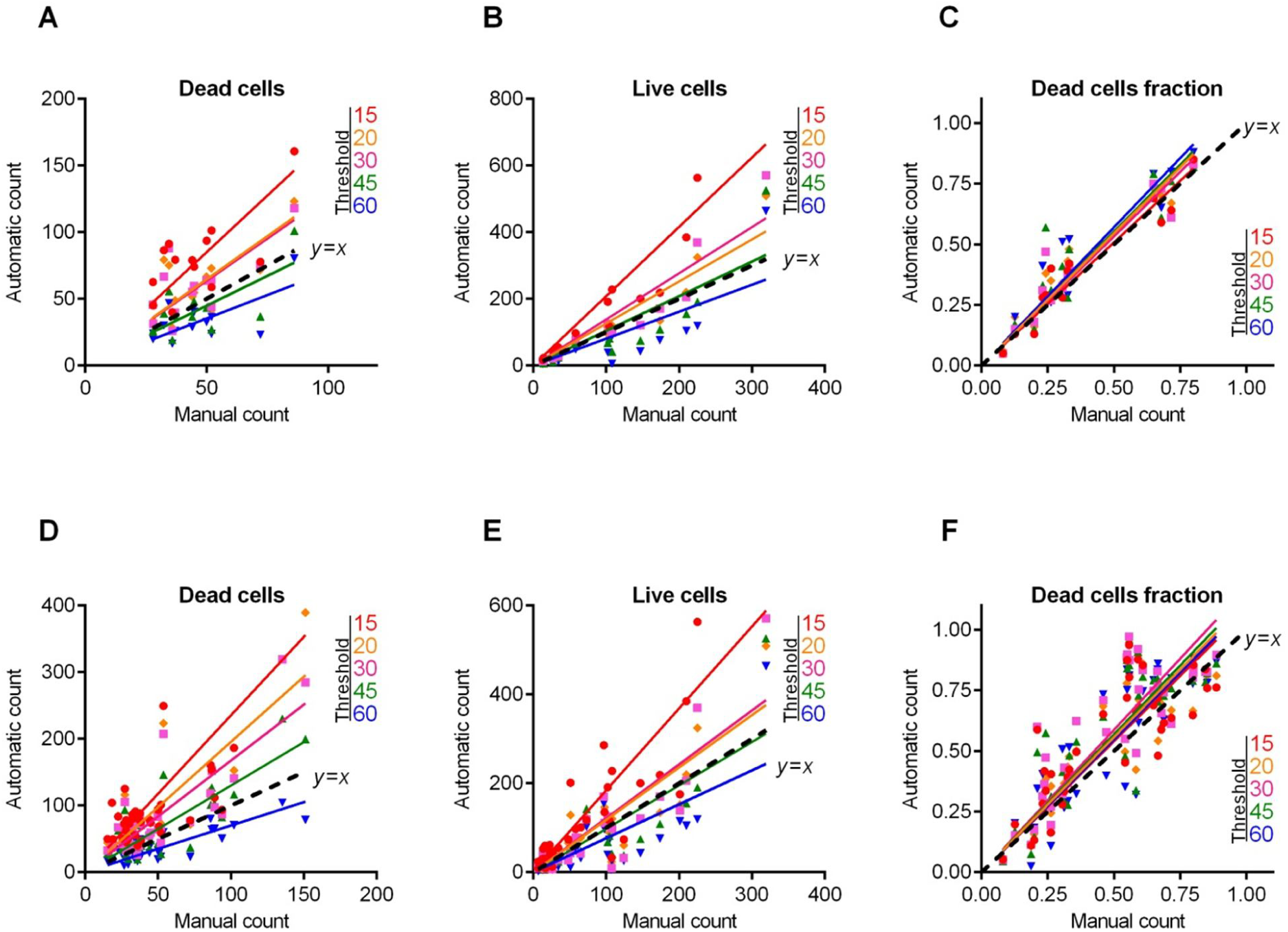
*B. subtilis* cells count and live/dead ratio dependence on the image analysis threshold *T*. Full lines show the linear regression lines, while dashed line shows the ideal counting line as determined by the manual analysis performed by several experts in visual microscopic image analysis. Panels A-C show data for the 12 images presented in Fig 4. Panels D-F show data for the 50 microscopic images taken randomly from different experiments with various imaging conditions.

**Table 2.** Regression coefficients *k*, the coefficients of determination *R*^2^ and the accuracy coefficient *L*^2^ indicating the correspondence between the automatic and the manual counting for the *B.subtilis* live/dead ratios on fluorescent images

| Threshold | 12 images |  |  | 50 images |  |  |
| --- | --- | --- | --- | --- | --- | --- |
| | $k$ | $R^2$ | $L^2$ | $k$ | $R^2$ | $L^2$ |
| 15 | 1,02 | 0,92 | 1,00 | 1,08 | 0,72 | 0,97 |
| 20 | 1,09 | 0,93 | 0,97 | 1,11 | 0,74 | 0,95 |
| 30 | 1,07 | 0,91 | 0,98 | 1,17 | 0,75 | 0,90 |
| 45 | 1,10 | 0,84 | 0,96 | 1,14 | 0,74 | 0,93 |
| 60 | 1,14 | 0,90 | 0,93 | 1,10 | 0,74 | 0,96 |

While the absolute cell count depends on the image analysis threshold which should be adjusted manually and therefore has a factor of subjectivity, the live/dead ratio quantification does not depend on the threshold settings in the range of 15-60 color intensity units (on the 0..255 scale) exhibiting linear function with *R*^2^ values exceeding 0.9 and regression coefficients of *k* ranging between 0.9 and 1.1 (Tables 1 and 2). This fact allows performing automatic analysis of multiple images with constant settings optimized from the first image in a series.

### Eukaryotic cells counting

The performance of *BioFilmAnalyzer* software was also analyzed for the eukaryotic cells. For that, Caco-2 cells were treated with different concentrations of camptothecin and analyzed after 24h of exposition. Figs 6 and 7C show the fractions of viable cells quantified with *BioFilmAnalyzer* software on microphotographs with overlapped green and red channels. Here, the best fit of manual and automatic count of either live or dead cells was observed for the analysis threshold *T* =30 (Fig 7). In contrast to bacterial cells, the accuracy was lower (*L*^2^ over 0.80), probably, because of the discrepancies in the cell sizes and uneven staining of cells. In contrast, when the software performance was evaluated on a series of 87 randomly chosen images (Figs 7D–7F), the automatic estimation of live/dead fractions fits with manual one with high confidence level (*L*^2^ equals 1.0, see Table 3). This effect could be attributed to larger effective sizes and thus also smaller average number of cells in each field of view for eukaryotic cells compared to bacterial cells that in turn requires analyzing more images in order to obtain similar statistics.

**Fig 6.**
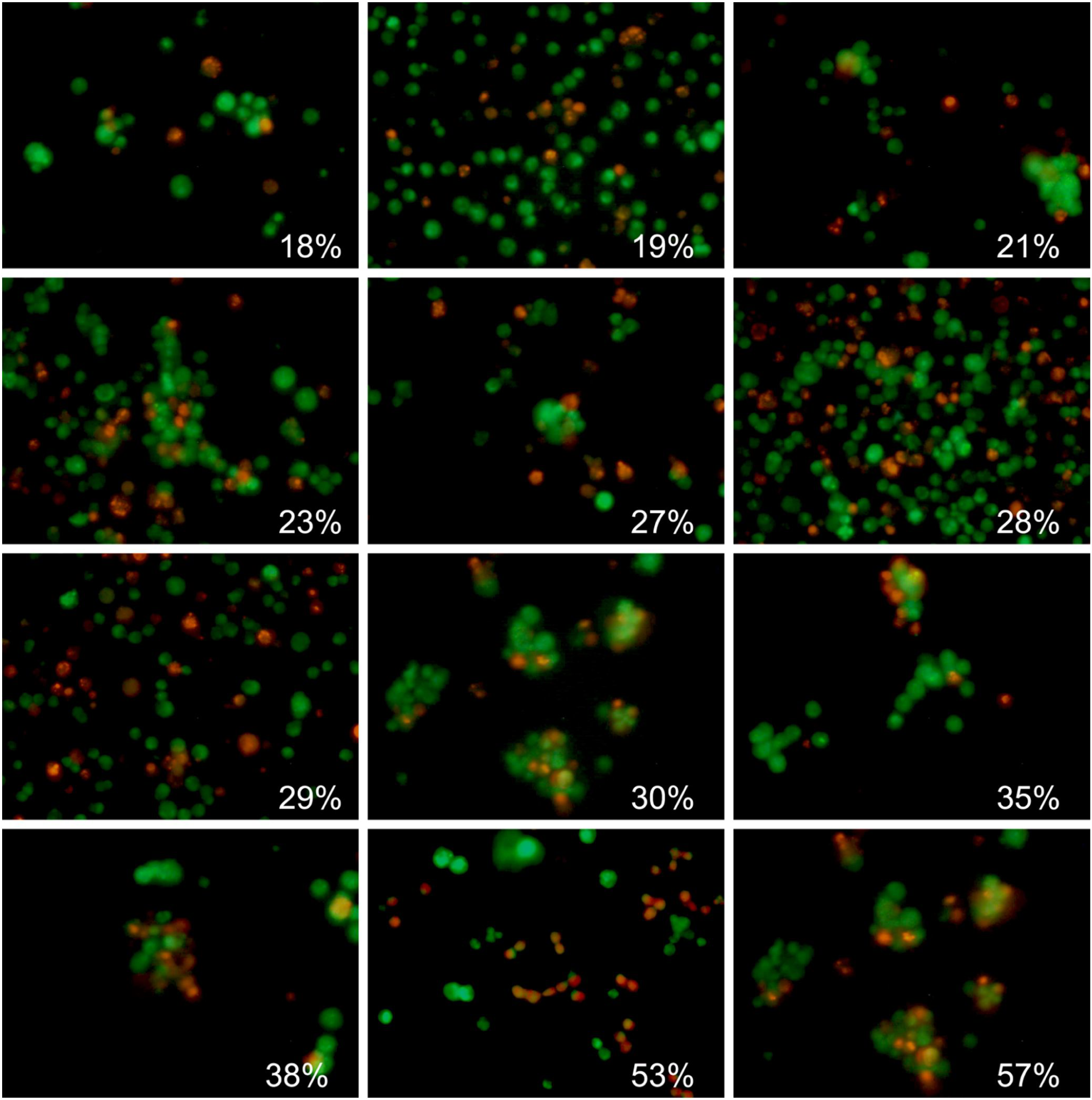
Evaluation of the Caco-2 non-viable cells fraction by using *BioFilmAnalyzer* software exemplified for 12 microscopic images. The percentage of red-stained cells quantified by the *BioFilmAnalyzer* software is shown in each panel

**Fig 7.**
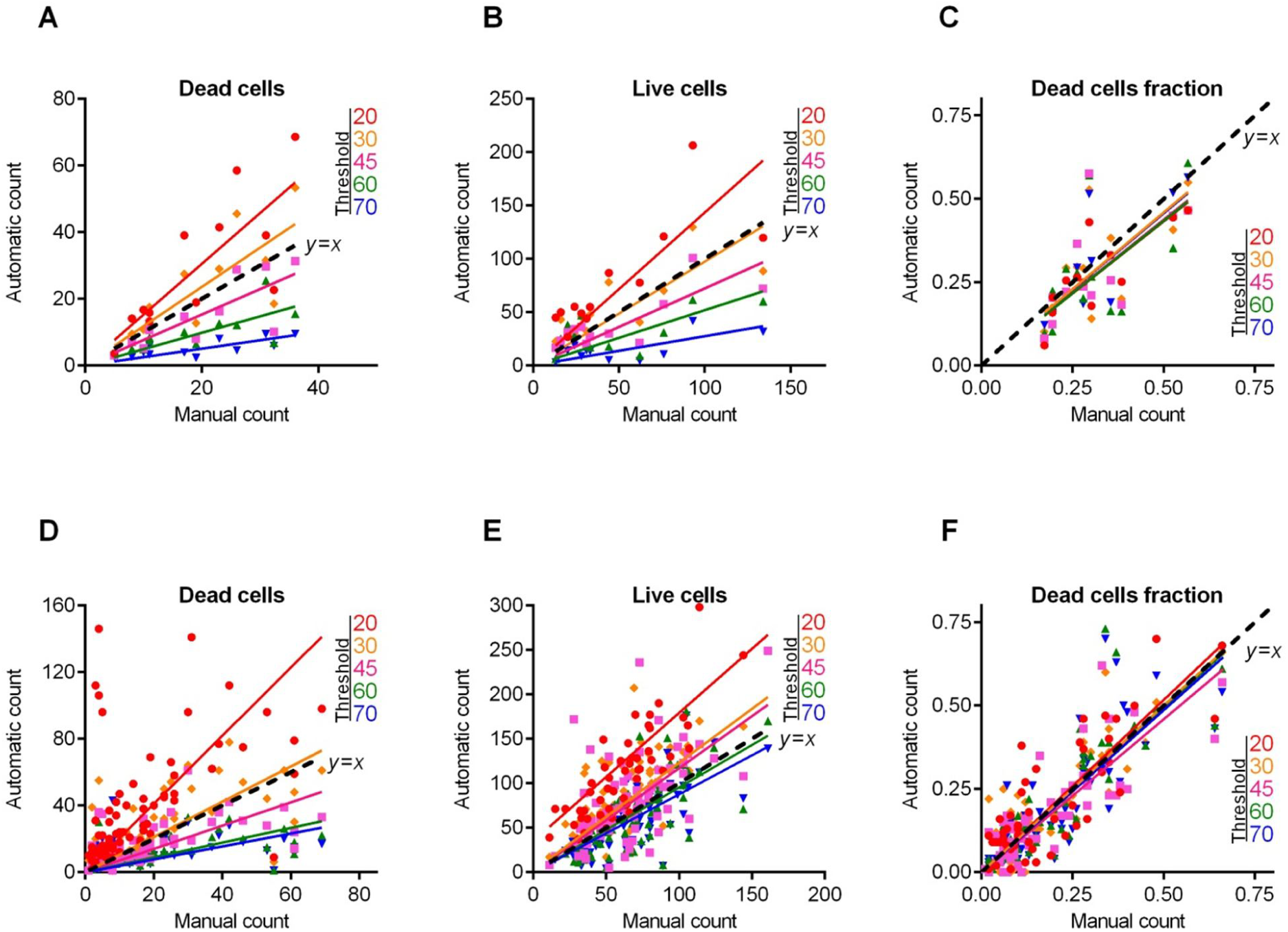
Caco-2 cells count and live/dead ratio dependence on the image analysis threshold *T*. Full lines show the linear regression lines, while dashed line shows the ideal counting line as determined by the manual analysis performed by several experts in visual microscopic image analysis. Panels A-C show data for the 12 images presented in Fig 6. Panels D-F show data for the 87 microscopic images taken randomly from different experiments with various imaging conditions

**Table 3.** Regression coefficients *k*, the coefficients of determination *R*^2^ and the accuracy coefficient *L*^2^ indicating the correspondence between the automatic and the manual counting for the Caco-2 live/dead ratios on fluorescent images

| Threshold | 12 images |  |  | 87 images |  |  |
| --- | --- | --- | --- | --- | --- | --- |
| | $k$ | $R^2$ | $L^2$ | $k$ | $R^2$ | $L^2$ |
| 15 | nd | nd | nd | nd | nd | nd |
| 20 | 0,86 | 0,69 | 0,80 | 1,03 | 0,73 | 1,00 |
| 30 | 0,92 | 0,53 | 0,93 | 1,00 | 0,76 | 1,00 |
| 45 | 0,88 | 0,46 | 0,83 | 0,92 | 0,71 | 0,97 |
| 60 | 0,87 | 0,40 | 0,81 | 0,99 | 0,73 | 1,00 |
| 70 | 0,91 | 0,54 | 0,92 | 0,97 | 0,71 | 1,00 |

To verify the accuracy of the overall procedure including both microscopic imaging and automatic cells counting, treated cells were detached from the wells by trypsin treatment, stained with DioC6 and ethidium bromide and aliquoted. One half of the sample was analyzed with flow cytometry to evaluate the fraction of necrotic cells, while the other half was subjected to microscopy and quantified by *BioFilmAnalyzer*. Fig 8 shows fractions of non-viable cells in 4 independent repeats quantified with flow cytometry versus automatic analysis of series of 10 microphotographs from each sample with separate green and red channels (representative examples of images are shown in the figure).

**Fig 8.**
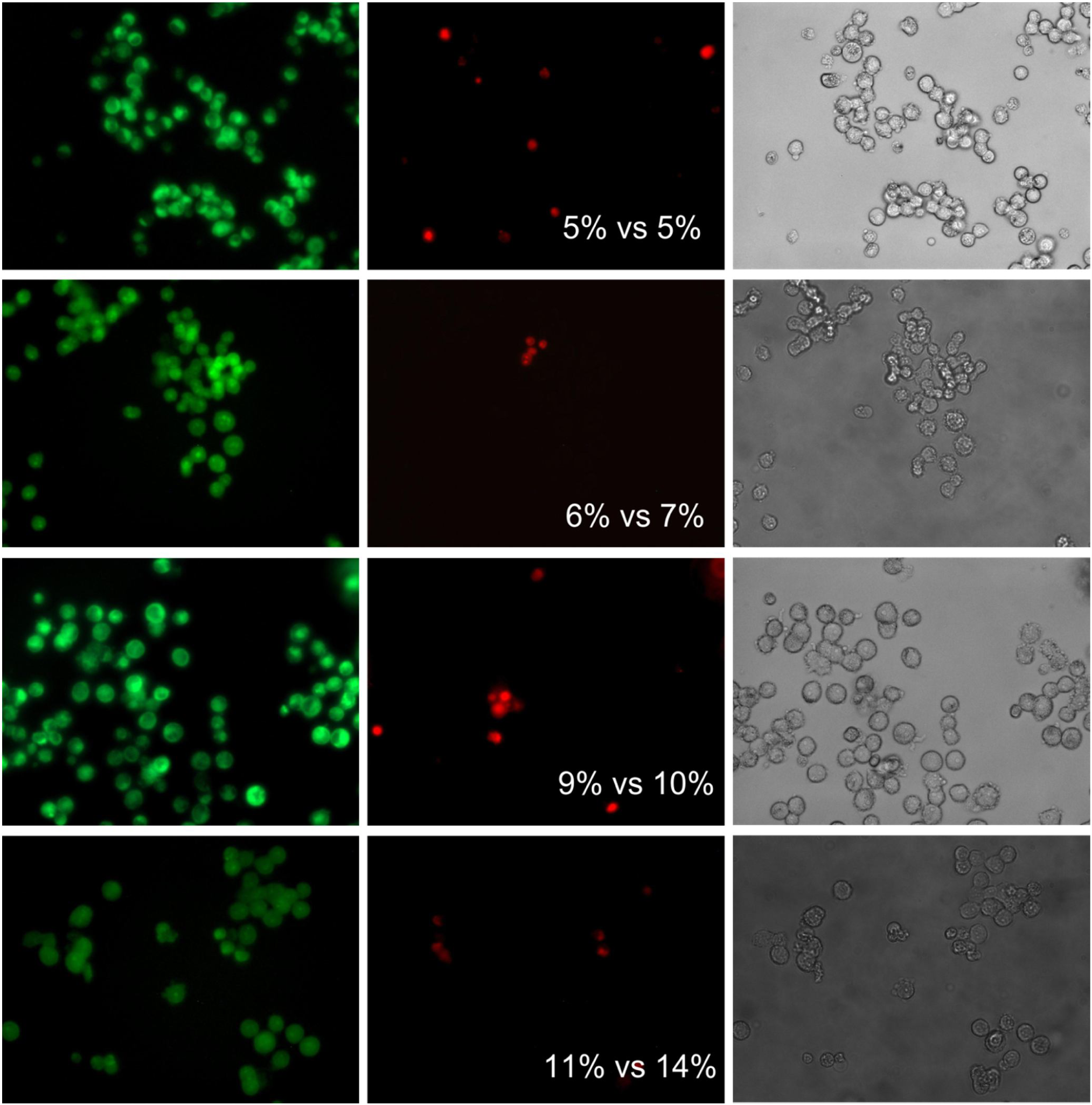
The fraction of the non-viable eukaryotic cells quantified by flow cytometry versus automatic analysis with *BioFilmAnalyzer*.

### Quantification of confocal images with Z-stacks

Finally, we used the same algorithm and software to analyze confocal images with Z-stacks, including 4 confocal images of *S. aureus* cells treated with different antimicrobials from our recent work [22, 23]. For that, we analyzed the raw series of 2D images that were used previously to reconstruct 3D images in [22, 23], where the fraction of live/dead cells were evaluated by both automatic and manual expert counting (Fig 9).

**Fig 9.**
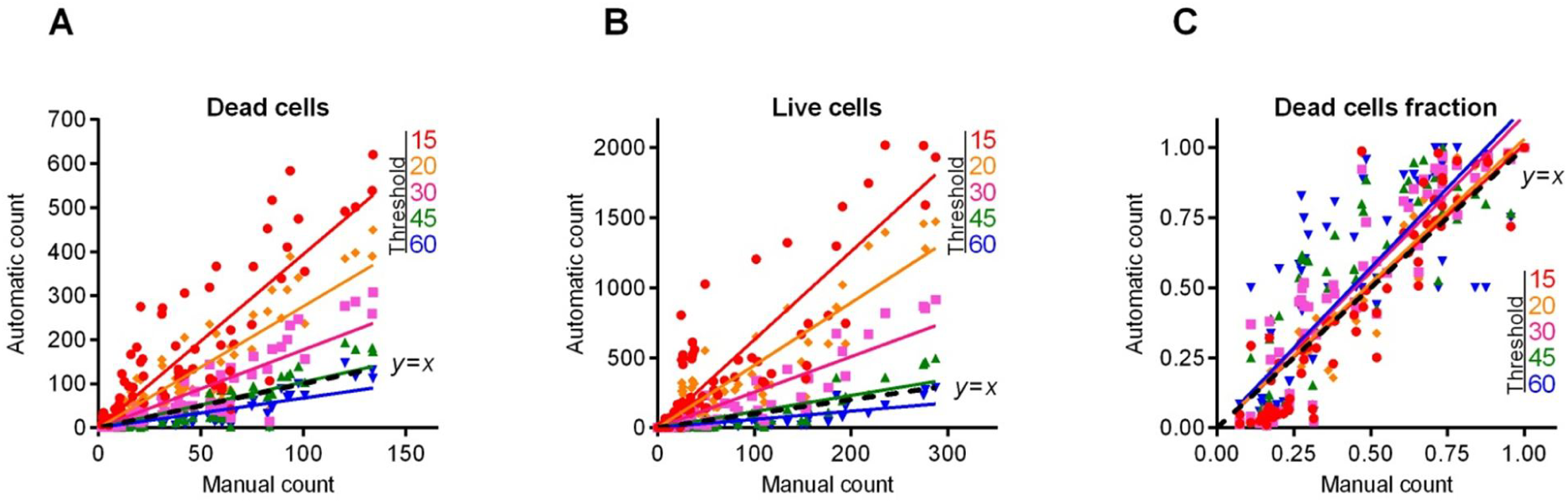
Automatic vs manual cell count cell count and live/dead ratio dependence for different image analysis thresholds *T*. Full lines show the linear regression lines, while dashed line shows the ideal counting line as determined by the manual analysis performed by several experts in visual microscopic image analysis. Four different confocal images containing between 12 and 23 Z-stacks each were taken for the analysis.

Typically 2d images obtained by confocal layer scanning technique are lower quality in comparison with single layer fluorescent microscopic images considered above. There are generally several contributing effects including defocus aberration of the cells that appear close but nevertheless displaced against the focal point due to the limited depth of focus as well as motion blur due to the sample shift during operation. Both effects lead to blurred images, the problem that is partially resolved in the 3d reconstruction algorithms by averaging or smoothening filters that improve the overall image quality at the cost of its effective resolution. The question is, whether raw 2d images obtained by confocal layer scanning technique can be used for the cell sub-population quantification using our algorithm, and whether this would require some preliminary filtering to reduce blurring effects. For the latter, two standard image filtering techniques, namely the Gaussian and the Sobel filters have been tested. The Sobel filtering aims on edge detection in the images using a discrete differentiation operator which computes an approximation of the gradient of the image intensity function. Therefore this kind of image preparation might be helpful to make edges of cells more stepwise and thus to reduce the dependence of the performance of the cell counting algorithm on the choice of the threshold that may then appear anywhere within this step. Alternatively, blurring may be treated as effective additive noise that could be reduced with simple Gaussian filter. Exhibiting a Gaussian impulse response, such filter decreases the overall noise level in the image.

Fig 10 shows an example of 2d confocal Z-stack microscopic image before and after Gaussian and Sobel filtering (upper panel) and the regression functions of automatic cells count at different thresholds as a function of the manual cell count (lower panel) before and after preliminary filtering of the images (obtained for the entire cohort of studied confocal images). The figure shows that, while the image appears visually less blurred, there is no significant improvement on cells fraction count according to the regression analysis results (see also Table 4). Furthermore, preliminary filtering leads to the overestimation of the red-stained cell count in some samples analyzed, this way also corrupting the overall sub-population fraction estimates. Therefore, we find that due to the general robustness of the sub-population fraction estimation against moderate variations of the threshold *T* (and thus also its relation with the quantile of the image color distribution), 2d images obtained by confocal layer scanning technique can be analyzed without preliminary image filtering.

**Fig 10.**
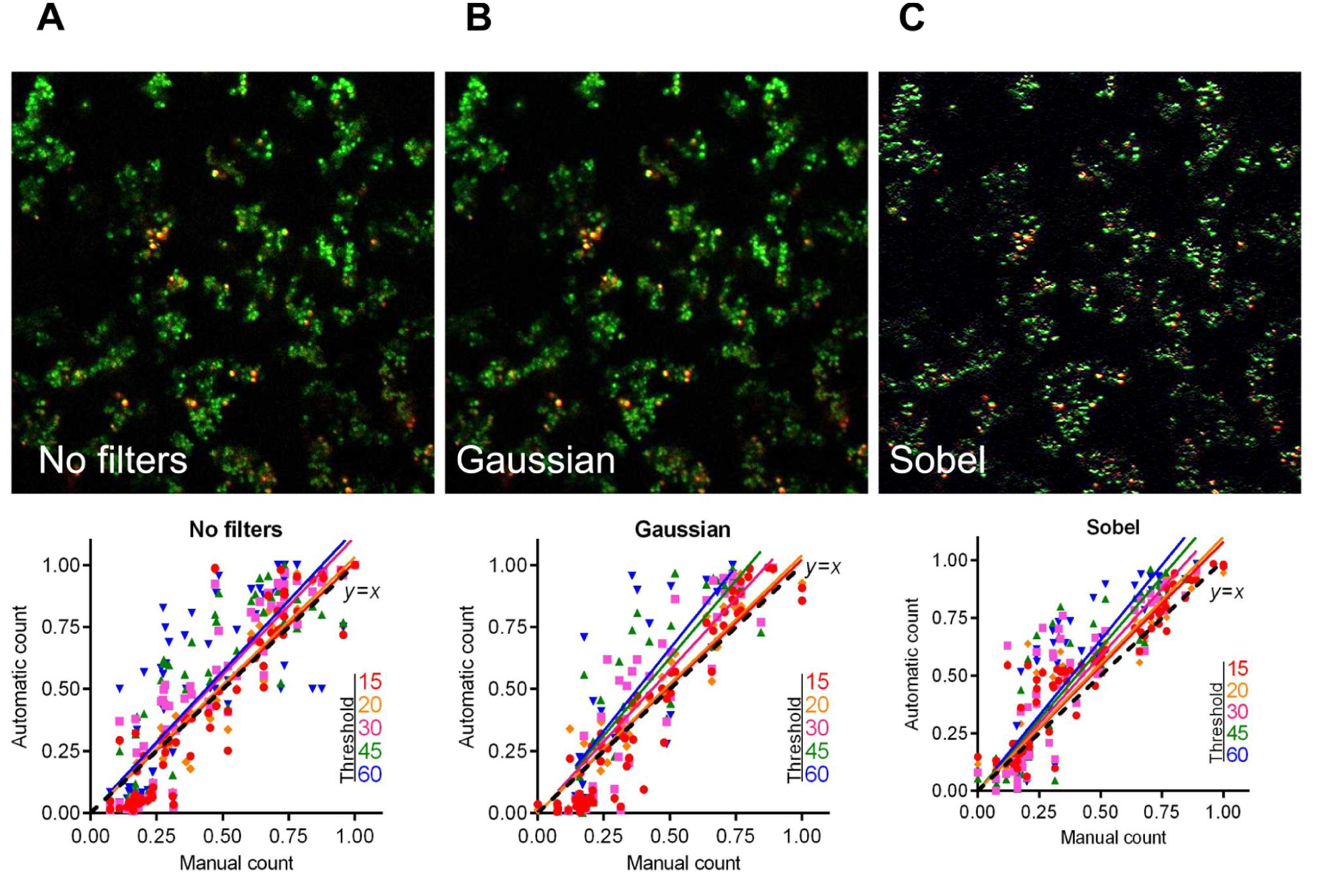
The live/dead ratio dependencies for different image analysis thresholds *T*. Full lines show the linear regression lines, while dashed line shows the ideal counting line as determined by the manual analysis performed by several experts in visual microscopic image analysis. Four confocal images containing between 12 and 23 Z-stacks were taken for the analysis, before and after being subjected to either Gaussian or Sobel filtering, as indicated.

**Table 4.** Regression coefficients *k*, the coefficients of determination *R*^2^ and the accuracy coefficient *L*^2^ indicating the correspondence between the automatic and the manual counting for the *S.aureus* live/dead ratios on confocal images.

|  | No filters |  |  | Gaussian |  |  | Sobel |  |  |
| --- | --- | --- | --- | --- | --- | --- | --- | --- | --- |
| Threshold | k | $R^2$ | $L^2$ | k | $R^2$ | $L^2$ | k | $R^2$ | $L^2$ |
| 15 | 1,01 | 0,81 | 1,00 | 1,02 | 0,85 | 1,00 | 1,08 | 0,87 | 0,98 |
| 20 | 1,03 | 0,81 | 1,00 | 1,04 | 0,83 | 0,99 | 1,10 | 0,84 | 0,97 |
| 30 | 1,11 | 0,82 | 0,95 | 1,15 | 0,76 | 0,92 | 1,17 | 0,77 | 0,92 |
| 45 | 1,14 | 0,77 | 0,93 | 1,25 | 0,71 | 0,78 | 1,24 | 0,79 | 0,87 |
| 60 | 1,14 | 0,68 | 0,93 | 1,32 | 0,60 | 0,69 | 1,30 | 0,78 | 0,78 |

## Adjustment of the threshold and single cell size balance

As we have already mentioned above, typical single cell sizes vary for different thresholds *T*, and thus the choice of these parameters should be properly balanced. Therefore, in general, for every chosen single cell size range there is an optimal threshold value *T*, and vice versa. Our results indicate that moderate threshold imbalance does not affect the overall statistics significantly, especially when averaging over large image series. However, in some cases adaptive threshold adjustment is nevertheless required, either for overcoming significant imbalance between different color channels or for the comparison between series of images obtained under considerably different staining and imaging conditions which are unavoidable, such as independent experimental repeats obtained in different time frames, with modified staining agent, different imaging settings, replaced lightning or optical components and so on.

To overcome this issue, we next suggest a simple adaptive threshold adjustment procedure that allows for an easy choice of the threshold value *T* for particular single cell sizes based on a single objective criterion. This procedure is performed once before running the analysis of a new image series given that the image parameters differ considerably from the previously analyzed one. The entire procedure is based on the multi-threshold analysis and calculation of empirical histograms of isolated objects sizes for every tested threshold value. These histograms are exemplified in Figs 11–13 for three representative microscopic images of both coccal and rod bacterial cell morphologies as well as for the eukaryotic cell lines, with raw images shown in panels A of respective figures. In all figures, panels B and C show the isolated objects belonging to a given size range after applying threshold *T* and cell size adjustment exemplified for the red and green color channels, respectively. In panel D, the upper (black) curve gives the total area above the threshold *T*, which equals the sum of the lower (colored) curves representing the total area of isolated objects within a given size range.

**Fig 11.**
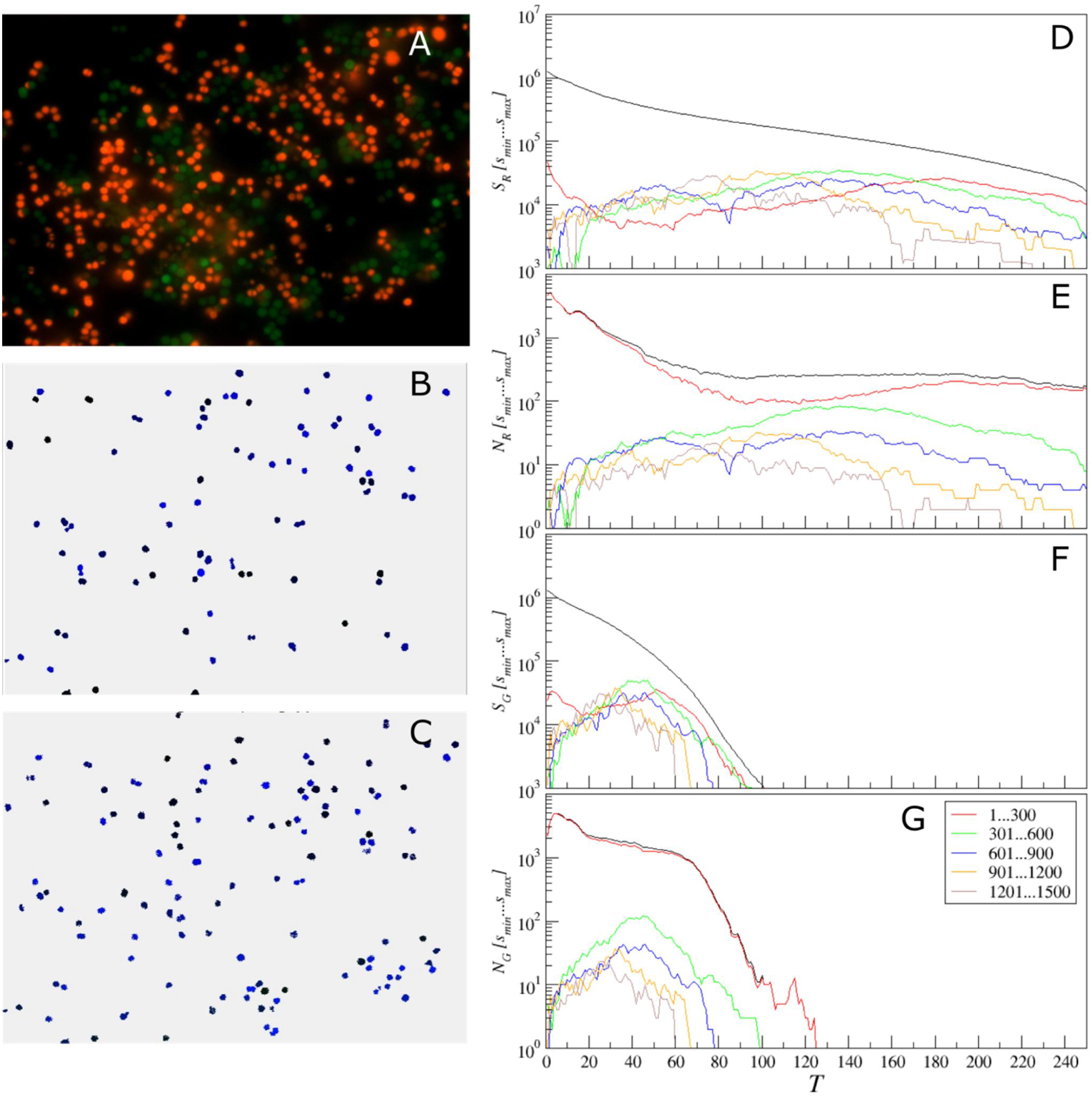
Adjustment of the threshold and single cell size balance for *S. aureus*. (A) - the raw image, (B) and (C) – isolated objects belonging to a given size range between 301 and 600 pixels, exemplified for the red (with *T* = 140) and the green (with *T* = 45) color channels, respectively. (D, F) the black curve gives the total area above the threshold *T*, which is the sum of the lower (colored) curves that represent the total area of isolated objects within a given size range for the red (D) or the green channel (F). (E, G) the black curve gives the total count of objects that appear isolated after threshold-based filtering for any given threshold value *T*, which is the sum of the lower (colored) curves that represent the total count of isolated objects within a given size range for the red (D) or the green channel (F).

**Fig 12:**
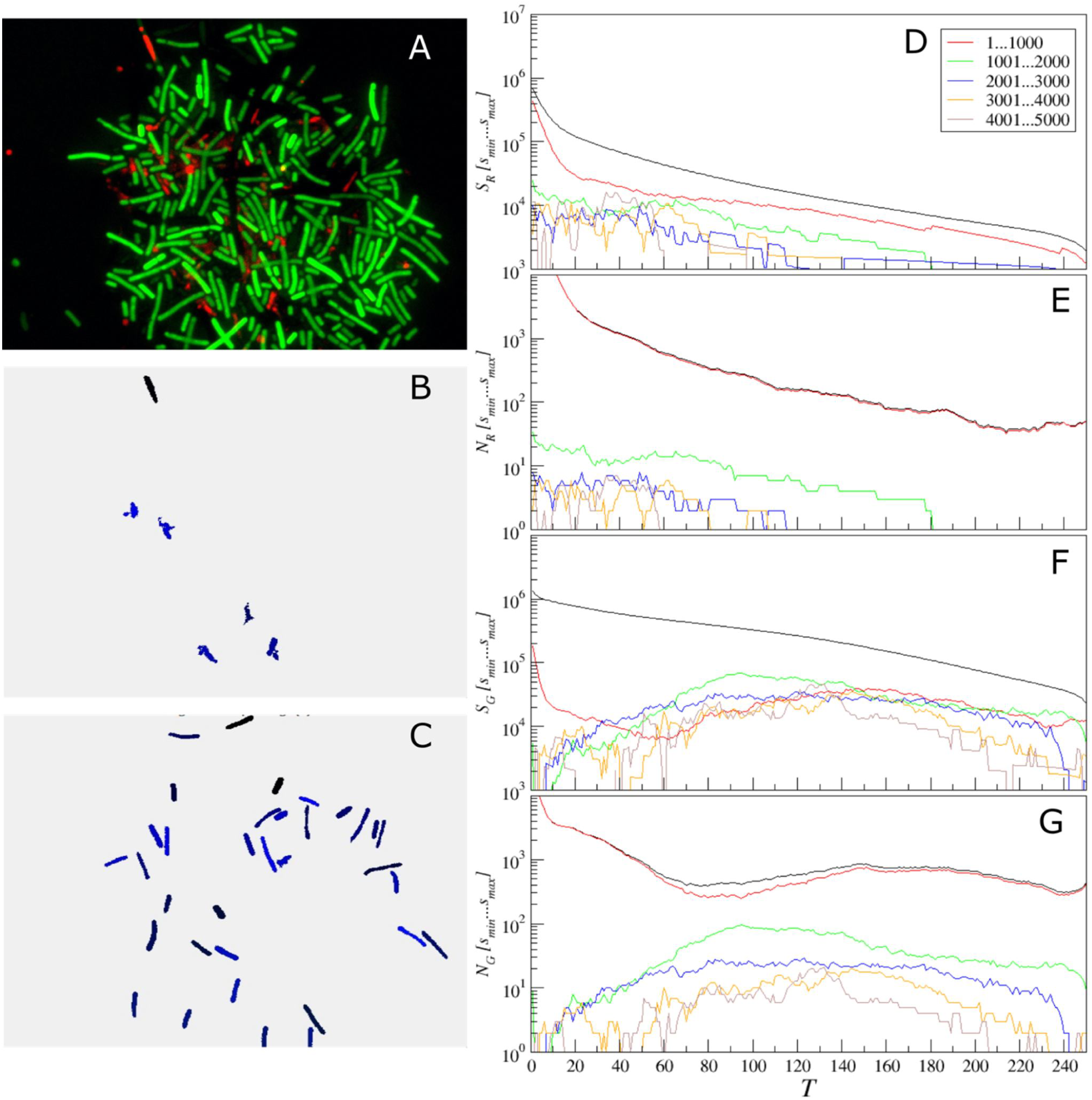
Adjustment of the threshold and single cell size balance for *B. subtilis*. (A) - the raw image, (B) and (C) – isolated objects belonging to a given size range between 1001 and 2000 pixels, exemplified for the red (with *T*= 70) and the green (with *T*= 100) color channels, respectively. (D, F) the black curve gives the total area above the threshold *T*, which is the sum of the lower (colored) curves that represent the total area of isolated objects within a given size range for the red (D) or the green channel (F). (E, G) the black curve gives the total count of objects that appear isolated after threshold-based filtering for any given threshold value *T*, which is the sum of the lower (colored) curves that represent the total count of isolated objects within a given size range for the red (D) or the green channel (F).

**Fig 13:**
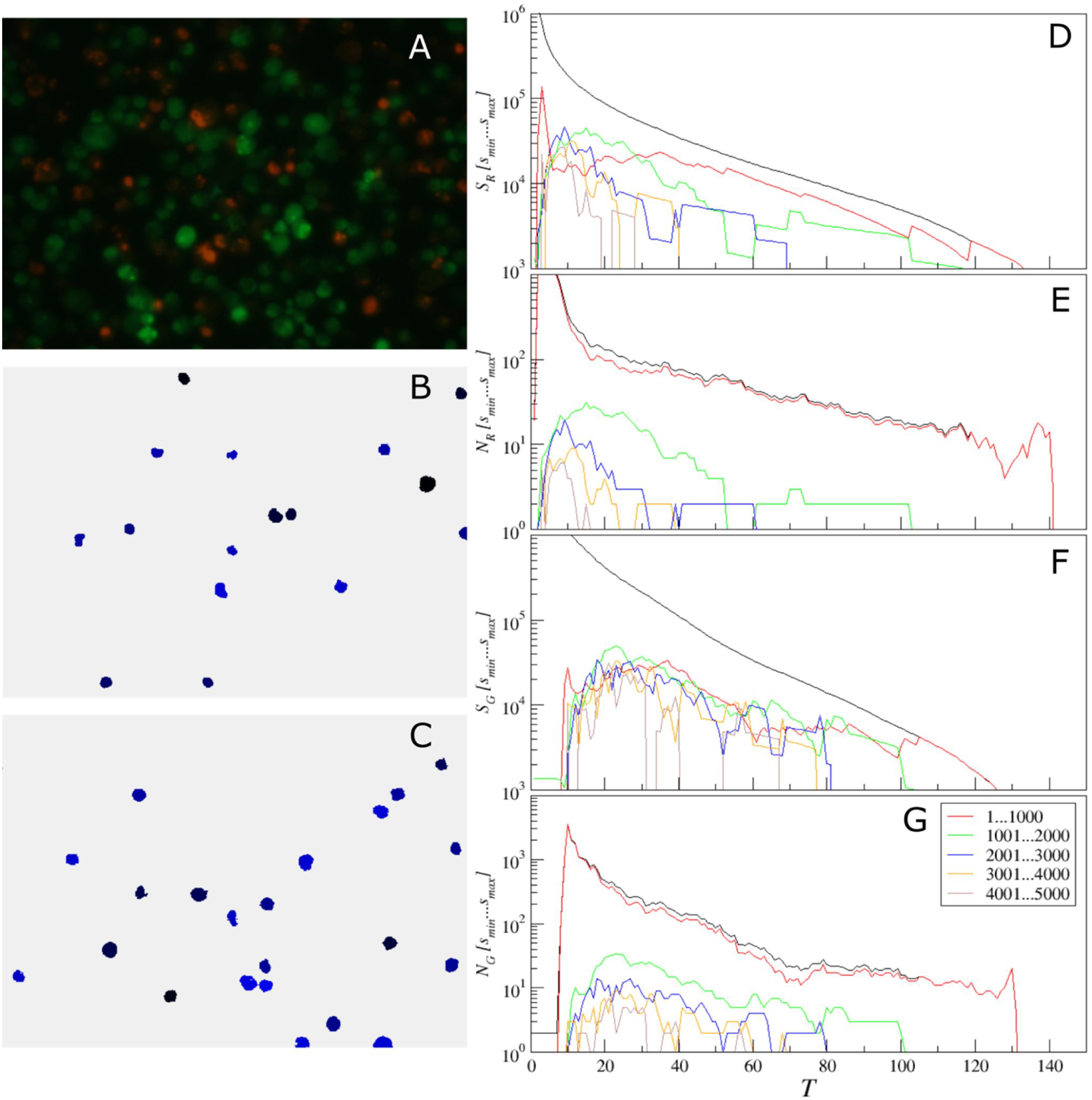
Adjustment of the threshold and single cell size balance for eukaryotic cells. (A) - the raw image, (B) and (C) – isolated objects belonging to a given size range between 1001 and 2000 pixels, exemplified for the red (with *T* = 15) and the green (with *T* = 25) color channels, respectively. (D, F) the black curve gives the total area above the threshold *T*, which is the sum of the lower (colored) curves that represent the total area of isolated objects within a given size range for the red (D) or the green channel (F). (E, G) the black curve gives the total count of objects that appear isolated after threshold-based filtering for any given threshold value *T*, which is the sum of the lower (colored) curves that represent the total count of isolated objects within a given size range for the red (D) or the green channel (F).

Similarly, in panel E the upper (black) curve gives the total count of objects that appear isolated after threshold-based filtering for any given threshold value *T*, which equals the sum of the lower (colored) curves representing the total count of isolated objects within a given size range. Similar quantities for the green color channel are shown in panels F and G, respectively. These histograms are in direct correspondence to the key quantities used in our algorithm, since the ratio of the respective color curves gives us the effective single cell size, while the black curve in panel D divided by this effective cell size gives us the effective cell count. By comparing panels D vs E as well as F vs G one can see that the histograms exhibit similar shapes, thus indicating that their ratio is relatively stable, again confirming the robustness of our algorithm against moderate threshold imbalance. Furthermore, when black curves for both color channels (red and green) depicted in panels D and F decrease with increasing threshold *T* by the same functional form, such that their ratio is roughly constant at least within a certain threshold range, *any* chosen threshold *T* within this range would, by definition, lead to the same relative fractions of counted cells. Moreover, for a series of images, it is sufficient that this condition is fulfilled for all analyzed images in average. This shows explicitly why in the previous tests we obtained similar results for relative quantities for a wide range of threshold values *T*, while individual cell counts were in some cases overestimated for small and underestimated for large thresholds *T*, in comparison with manual count. Prominent exceptions from this rule can be observed in Fig 11, where pronounced color imbalance can be observed. In contrast, Figs 12 and 13 show that even under moderate color imbalance the above conditions are met to a certain extent.

The figures also show that the overwhelming majority of isolated objects are small objects, which are below 300 pixels for coccal cells and below 1000 pixels for both rod and eukaryotic cells, respectively. Let us next focus on Figs 11D and 11E. In a marked contrast to the red curve for the small objects, histograms for larger sizes exhibit characteristic maxima, which are most pronounced in the green curve for the objects of sizes between 301 and 600 pixels. Assuming that the most common objects of similar size in the image except small noise bursts are single cells, we can choose the optimized threshold from the respective histogram maximum, which is around *T* = 140. For larger sizes, the blue histogram splits into two characteristic regimes with local maxima at around *T* = 55 and 130, respectively. This shows strong non-homogeneity of the objects that belong to this size that is untypical for single coccal cells, indicating that these are likely cell clusters. For larger sizes, the histograms of object counts are well below the green curve, indicating that objects of these sizes are less common than typical single cells, and significantly fluctuating due to the lack of statistics. Thus we next set the threshold T = 140 and size range 301…600 and obtain the results of single cell selection depicted in panel B of Fig 11.

Now similar procedure can be easily repeated for the green channel. As one can see clearly from Figs 11F and 11G, while the green curve is again above the curves for larger sizes, due to the considerably lower intensity of the green channel, the optimal threshold values is now around *T* = 45. After the corresponding threshold adjustment, we obtain the typical examples of single cells depicted in Fig 11C.

Similar conclusions can be drawn from the analysis of Figs 12 and 13 for rod and eukaryotic cells, respectively. In both cases, following the same criterion based on finding the maximum of the histogram for the most common objects leads to reasonable selection of typical single cells, as indicated by panels B and C for the red and green channels, respectively.

Therefore, the suggested approach allows dealing with images obtained with considerable color imbalance as well as comparison of repeats obtained under different staining and imaging conditions that are sometimes unavoidable in routine lab practice, due to equipment and reagent aging, repairs, and replacements. Since all further analysis is based on the effective number of cells that is determined independently for each color channel with its own optimized threshold and single cell size values, the above adjustments do not affect any further stages of analysis.

## Conclusion

To summarize, our results indicate that the sub-population fraction estimates obtained from fluorescent-stained cells imaging data by *BioFilmAnalyzer* are very close to the results obtained by other techniques such as expert manual counting and flow cytometry. The two-step algorithm implemented in the *BioFilmAnalyzer* that normalizes the stained image area in the units of the effective single cell size that are determined following the suggested objective criterion while remaining under manual consistency control by the investigator performs largely independently of the cell shape and imaging conditions providing feasible results for cells aggregated in clusters. Moreover, it requires neither preliminary preparation nor filtering of raw images.

Thus, we suggest that the proposed algorithm implemented as the *BioFilmAnalyzer* software can be used for numerous applications. First, preliminary evaluation of cell counts and live/dead ratios can be quickly obtained without expertise in image processing. Second, the analysis of surface-adherent bacterial or eukaryotic cells without their resuspension and maintenance of the native distribution pattern on the surface is possible. Third, the quantification of cellular sub-populations from 2d confocal layer images is also possible. Fourth, the software allows quantification of the cells sub-populations expressing fluorescent proteins during long-term incubation. Finally, because no further user intervention is required after few initial adjustment and cross-check procedures which usually take a couple of minutes using few representative images, further processing can be done over coffee-break by automated analysis of a series consisting up to 100 images.

We believe that the suggested algorithm and software would be useful in saving time and efforts in cells sub-population quantification from fluorescent microscopy data that is commonly required in various biomedical, biotechnological and pharmacological studies, especially for the analysis of biofilm-embedded cells where the performance of conventional techniques based on cell counting or flow cytometry is strongly limited. Both the algorithm and the *BioFilmAnalyzer* can be freely downloaded at https://bitbucket.org/rogex/biofilmanalyzer/downloads/, utilized and redistributed.

## Supporting information

### BioFilmAnalyser.exe. Software module for Win32 (2000/XP and higher)

A software tool for semiautomatic processing of fluorescent microscopic images especially designed to overcome typical limitations of standard image analysis software under biofilm research conditions. Can be downloaded from https://bitbucket.org/rogex/biofilmanalyzer/downloads/

### BioFilmAnalyzer_User_Manual.pdf. The user manual

Can be downloaded from https://bitbucket.org/rogex/biofilmanalyzer/downloads/

